# Leaf shedding as a bacterial defense in Arabidopsis cauline leaves

**DOI:** 10.1101/189720

**Authors:** O. Rahul Patharkar, Walter Gassmann, John C. Walker

## Abstract

Plants utilize an innate immune system to protect themselves from disease. While many molecular components of plant innate immunity resemble the innate immunity of animals, plants also have evolved a number of truly unique defense mechanisms, particularly at the physiological level. Plant’s flexible developmental program allows them the unique ability to simply produce new organs as needed, affording them the ability to replace damaged organs. Here we develop a system to study pathogen-triggered leaf abscission in Arabidopsis. Cauline leaves infected with the bacterial pathogen *Pseudomonas syringae* abscise as part of the defense mechanism. *Pseudomonas syringae* lacking a functional type III secretion system fail to elicit an abscission response, suggesting that the abscission response is a novel form of immunity triggered by effectors. *HAESA/HAESA-like 2, INFLORESCENCE DEFICIENT IN ABSCISSION*, and *NEVERSHED* are all required for pathogen-triggered abscission to occur. Additionally *phytoalexin deficient 4, enhanced disease susceptibility 1, salicylic acid induction deficient 2*, and *senescence-associated gene 101* plants with mutations in genes necessary for bacterial defense and salicylic acid signaling, and NahG transgenic plants with low levels of salicylic acid fail to abscise cauline leaves normally. Bacteria that physically contact abscission zones trigger a strong abscission response; however, long distance signals are also sent from distal infected tissue to the abscission zone, alerting the abscission zone of looming danger. We propose a threshold model regulating cauline leaf defense where minor infections are handled by limiting bacterial growth, but when an infection is deemed out of control, cauline leaves are shed. Together with previous results our findings suggest that salicylic acid may regulate both pathogen‐ and drought-triggered leaf abscission.

**Author Summary:** Plants have a flexible development program that determine their form. We describe an organ level defense response in Arabidopsis to bacterial attack where plants simply shed heavily infected leaves. The genetics regulating this defense mechanism are comprised of both classical defense genes and floral organ abscission genes working together. Long distance signals are transmitted from infected areas to abscission zones which activate the abscission receptor. Salicylic acid, a defense hormone, signaling is necessary for cauline leaf abscission.

## Introduction

An arms race has been waged for eons between plants and microbial pathogens. Plants have evolved sophisticated defense mechanisms against disease while pathogens have acquired equally sophisticated means of avoiding the host’s defense. Plants lack an adaptive immune system and thus rely on an innate immune system to limit undesirable microbial colonization [1–3]. The plant innate immune system can detect microbial pathogens directly by recognizing microbe associated molecular patterns (MAMPS) that are bound by pattern recognition receptors (PRR) on the host cells [2,3]. Additionally, plants can scan themselves for general damage or modification caused by microbial pathogens, such as degradation of the plant cell wall that releases so-called damage-associated molecular patterns. Collectively, this part of the plant innate immune system is called pattern-triggered immunity (PTI) [2]. A second layer of plant immunity, effector-triggered immunity (ETI), relies on resistance proteins to detect pathogen effectors that pathogens deploy in the host cell to manipulate immune responses or release of nutrients [2,3]. Most commonly, these resistance proteins either directly bind specific effectors or detect effector-induced changes to host proteins with which they associate [1–4]. Both PTI and ETI have been well studied in Arabidopsis rosette leaves before flowering has occurred [1–7]. However, the Arabidopsis immune response is less understood in other tissues and at other developmental time points. Additionally, defense studies in plants have focused largely on microbe growth suppression and containment mechanisms at the tissue level but not the organ level.

Recently, we discovered that when Arabidopsis protects itself against drought by abscising its cauline leaves, it uses the same set of signaling components as are required for the shedding of flower petals after fertilization [8]. Cauline leaves are the aerial leaves attached directly to the inflorescence stem without a petiole (Supplemental Figure 1). Despite stark differences in the organs being abscised and the physiological and developmental basis that triggers abscission, signaling within abscission zones appears to be highly conserved. Cauline leaves and floral organs both require the redundant abscission receptor-like protein kinases HAESA and HAESA-like 2 (HAE/HSL2) which are triggered by a peptide derived from INFLORESCENCE DEFICIENT IN ABSCISSION (IDA). HAE/HSL2 activates a MITOGEN ACTIVATED PROTEIN KINASE (MAPK) cascade that in turn de-represses the MADS domain transcription factor AGAMOUS-LIKE-15 which in turn allows *HAE* to be expressed [8–11]. Newly produced HAE is then thought to be shuttled to the plasma membrane with the assistance of the ADP-ribosylation factor GTPase-activating protein NEVERSHED (NEV), which completes a positive feedback loop. Recent advances have revealed that the abscission receptor, HAE, utilizes SOMATIC EMBRYOGENESIS RECEPTOR-LIKE KINASES (SERK) as a co-receptor in the recognition of IDA peptide [12,13]. A recent breakthrough showed a 14mer biologically active IDA peptide is released from the IDA protein via the activity of subtilisin-like serine proteinases [14]. Interestingly, pieces of the abscission signaling pathway have been reported to be used by pathogenic bacteria to degrade pectin in rosette leaves and ease their colonization of the leaves [15].

While there is plentiful molecular and physiological knowledge detailing how Arabidopsis rosette leaves respond to a variety of pathogens, much less is known about how infected organs are shed to physically remove the attacker. What is known is that several plant species have documented abscission in response to disease. For example, tomato plants have been reported to shed their leaves in response to being vacuum infiltrated or dipped with *Pseudomonas syringae* [16,17]. Pepper plants shed leaves infected with *Xamthomonas campestris* pv. vesicatoria [18]. Powdery mildew has also been reported to cause leaf abscission in tomato [19]. Tomato Yellow Leaf Curl Virus can make tomato flowers abscise [20]. The fungus *Cercospora arachidicola* Hori triggers leaflet abscission in peanut [21]. Ethylene typically regulates the disease-triggered abscission [16,18,21]. The significance of shedding diseased organs at the plant level is that it enables plants to greatly reduce the titer of pathogens on the plant body. Bacterial pathogens like *Pseudomonas syringae* are spread by raindrop momentum, wind, and insects [22–25]. In contrast to microbial growth reduction mechanisms, shedding infected leaves allows plants to completely eliminate disease sources that may spread to healthy tissue.

We sought to further understand how plants contain bacterial infection by shedding entire affected organs. Therefore, we developed a *Pseudomonas syringae* pv. tomato strain DC3000 (DC3000) triggered cauline leaf system in Arabidopisis to understand plant defense mechanisms in leaves that can be shed. This system builds on the extremely well studied pathosystem in which rosette leaves from non-flowering Arabidopsis, accession Columbia are infected with *Pseudomonas syringae* pv. tomato strain DC3000 [26]. This work also builds on our previous work showing that Arabidopsis cauline leaves from flowering plants can be shed in response to drought [8]. This study shows that Arabidopsis uses leaf abscission as a bona fide defense mechanism against bacterial infection. The abscission response is robust and can be triggered by bacteria causing either disease or ETI. The abscission signal pathway originally elucidated in floral organs is necessary for the shedding defense response. Additionally, the defense hormone salicylic acid (SA) likely regulates leaf shedding since a number of genes necessary for SA mediated defense are also necessary for the full leaf shedding defense response.

## Results

### *HAE* is co-expressed with EDS1 and PAD4 in abscission zones

Co-expression analysis of *HAE* expression across many tissues and treatments revealed that *HAE* was co-expressed with *PHYTOALEXIN DEFICIENT 4* (*PAD4*) and *ENHANCED DISEASE SUSCEPTIBILITY 1* (*EDS1*) (Figure 1). Both *PAD4* and *EDS1* are statistically increased through the process of stamen abscission along with *HAE* (Figure 1) [27]. Interestingly, *HAE* has altered expression in tissue that has altered levels of SA. For example, *salicylic acid induction deficient 2* (*sid2*) mutants and NahG (salicylate hydroxylase) over-expressing plants have reduced levels of SA and also reduced *HAE* expression [28,29]. Conversely, *mpk4* and *mkk1 mkk2* mutant plants have increased levels of SA and also increased *HAE* expression (Figure 1) [30,31]. SA is a key hormone regulating the defense response and also the senescence process. Previously, it was shown that floral receptacles from *hae hsl2* mutant plants have altered expression of defense genes [32]. This wealth of circumstantial evidence suggested that defense and abscission may be connected.

**Figure 1.**
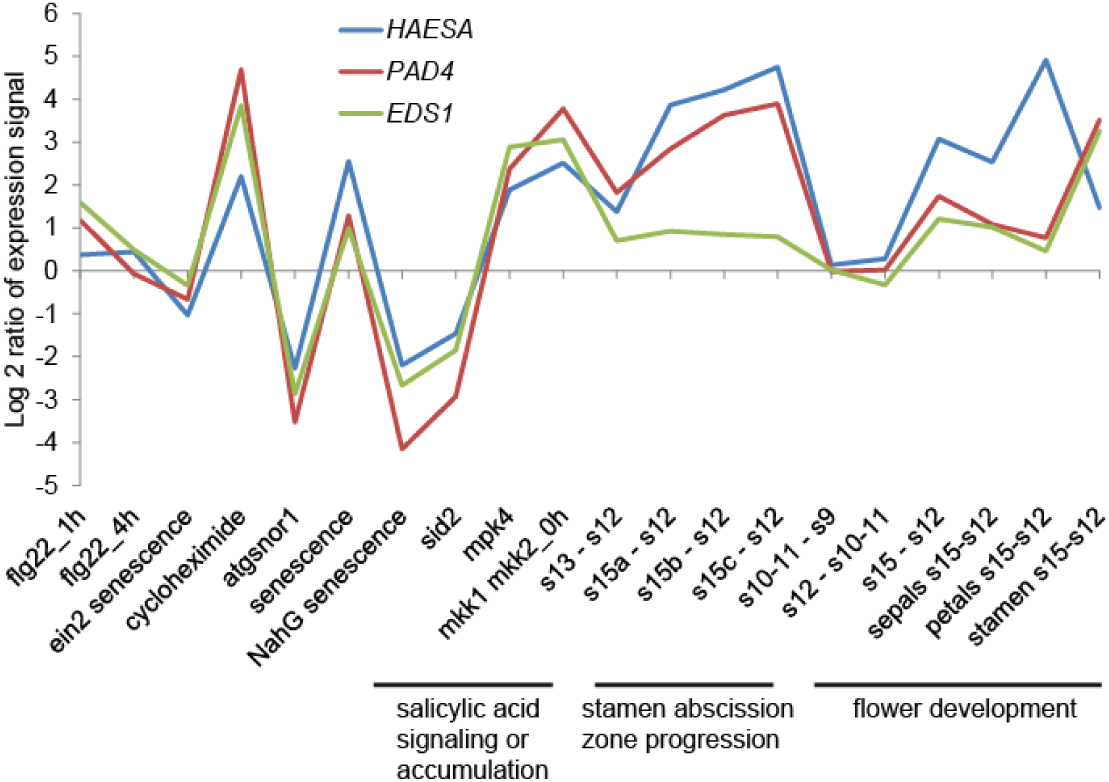
*HAESA* is co-expressed with *PAD4* and *EDS1* in a number of different tissues and treatments. Publicly available microarray data indicates that *PAD4* and *EDS1* are statistically increased during the abscission process in stamen abscission zones [27]. Furthermore, *HAE, PAD4*, and *EDS1* expression are up in shoot and leaf tissues of mutants with increased SA levels and down in mutants with decreased SA levels [49,50,31,51,52].

### DC3000 activates HAE and triggers cauline leaf abscission provided it has a functional Type III secretion system

Based on the gene expression data we wanted to test whether disease might also trigger cauline leaf abscission and if so understand how this works. Treatment of Columbia-0 (Col-0) cauline leaves with DC3000 resulted in a clear induction of HAE-YPF expression (driven by the native promoter) (Figure 2A). DC3000 carrying the effector genes *avrRps4* or *avrRpm1*, which Col-0 responds with ETI to, also induce HAE expression, while bacteria with the *hrcC* mutation that lack a functional Type III secretion system do not trigger HAE expression (Figure 2A). Virulent or ETI-eliciting bacteria with a functional Type III secretion system trigger cauline leaf abscission 3 days after infection while DC3000 *hrcC*^−^ does not (Figure 2B). These results indicate the mere presence of MAMPs is not sufficient to activate abscission, rather the abscission response requires the Type III secretion system. Interestingly, DC3000 with or without *avrRpm1* or *avrRps4* cause similar levels of abscission, which suggests an endogenous effector in DC3000 that does not elicit ETI in Col-0 is triggering cauline leaf abscission. The hypersensitive response is a mechanism used by plants to restrict bacterial growth by triggering a rapid plant cell death in the area surrounding the infection [33]. The hypersensitive response in cauline leaves functions in similar fashion as it does in rosette leaves, where *DC3000*(*avrRps4*) triggers an HR-less ETI. Only *DC3000*(*avrRpm1*) causes leaves to collapse within 20 hours of infiltration while DC3000 alone or with *avrRps4* do not (Figure 2C). Bacterial growth is also limited by recognized effectors in cauline leaves, however, leaves from flowering plants, grown in conditions suitable for reproduction, are more resistant to *Pst* than typical rosette leaves used for the *Pst*-Arabidopsis pathosystem (Supplemental Figure 4) [26]. Bacteria multiply to about 100 times higher levels in the classical Pseudomonas-Arabidopsis pathosystem than in our cauline leaf system (Supplemental Figure 4). Additionally, the bacterial phytotoxin coronatine, that partially mimics jasmonic acid, is not necessary to trigger leaf abscission (Figure 2D-E) [34]. This indicates it is more likely an effector injected through the Type III secretion system causes leaf abscission rather than an effect of coronatine.

**Figure 2.**
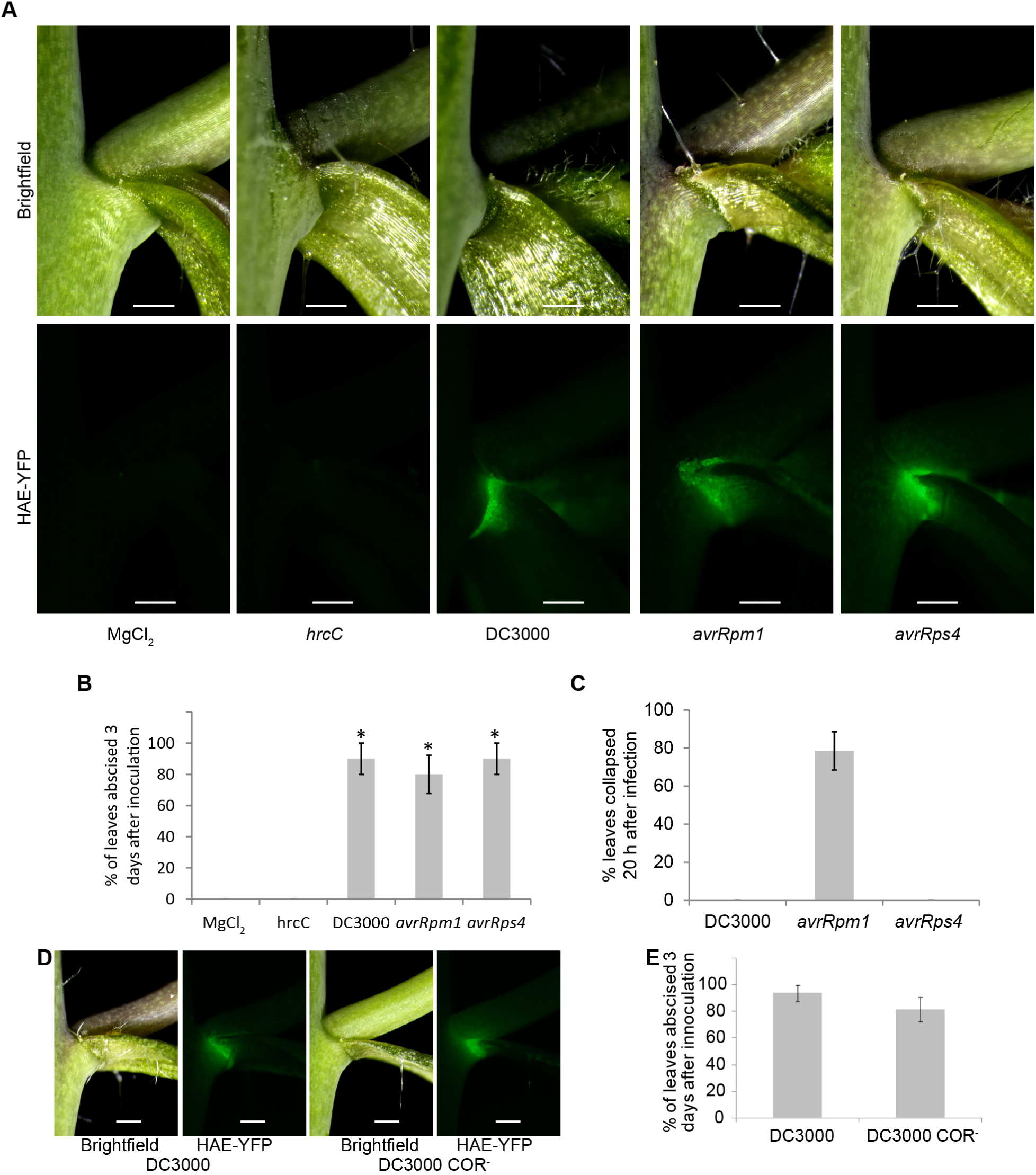
Bacteria with a viable type III secretion system can activate HAE expression and trigger cauline leaf abscission. (A) Plants expressing HAE-YFP (driven by the *HAE* promoter) were infected with virulent or avirulent DC3000. Images are 2 days after infiltration. The same samples are shown in the top and bottom panels where the top panels are imaged with reflected white light and the bottom panels are imaging YFP florescence. (B) Abscission of cauline leaves three days after infection. Data are mean ± s.e.m.; n=5 biological replicates (one plant each); t-test versus MgCl_2_ control; \**P* < 0.005. (C) Hypersensitive response in cauline leaves 20 h after infection. Data are mean ± s.e.m.; n=7 biological replicates (one plant each, 2 leaves per plant). (D) Leaves infected with DC3000 or DC3000 that does not produce coronatine (COR^-^) for two days. (E) Abscission of cauline leaves 3 days after infection. Data are mean ± s.e.m.; n=8 biological replicates (one plant each). Scale bar is 0.5 mm.

### Full abscission occurs when bacteria physically contact the cauline leaf abscission zone

To understand physiological mechanisms behind bacteria triggered abscission, we designed experiments to see whether the position of bacterial infiltration affected abscission. DC3000 was infiltrated so that it filled either the entire leaf, the proximal half of the leaf (closest to the AZ), the distal half of the leaf (away from the AZ), or the proximal quarter of the leaf (only on one side of the midrib). All leaf infiltration positions caused HAE-YFP to be expressed. However, bacteria that touched the AZ produced the strongest induction of HAE-YFP (Figure 3A, 3C). For example, in the quarter leaf infiltration, the side of the leaf with the bacteria produced stronger HAE-YFP than did the side not infiltrated (Figure 3A, 3C). Additionally, only infiltrations that touched the AZ resulted in full abscission (leaf falling off rather than being fully or partially attached) (Figure 3B). Again the extreme example of this is demonstrated by the quarter infiltration only triggering abscission on the side of the midrib where bacteria were present (Figure 3A, 3B, Supplemental Figure 2). Infiltrating the distal half of the leaf did cause partial abscission characterized by swelling of the AZ cells and some cell separation as well as expression of HAE-YFP (Figure 3 labeled “half away”, Supplemental Figure 3). This suggests that some signal is being transduced across the uninfected half of the leaf that affects the AZ or that bacteria move through the leaf toward the AZ.

**Figure 3.**
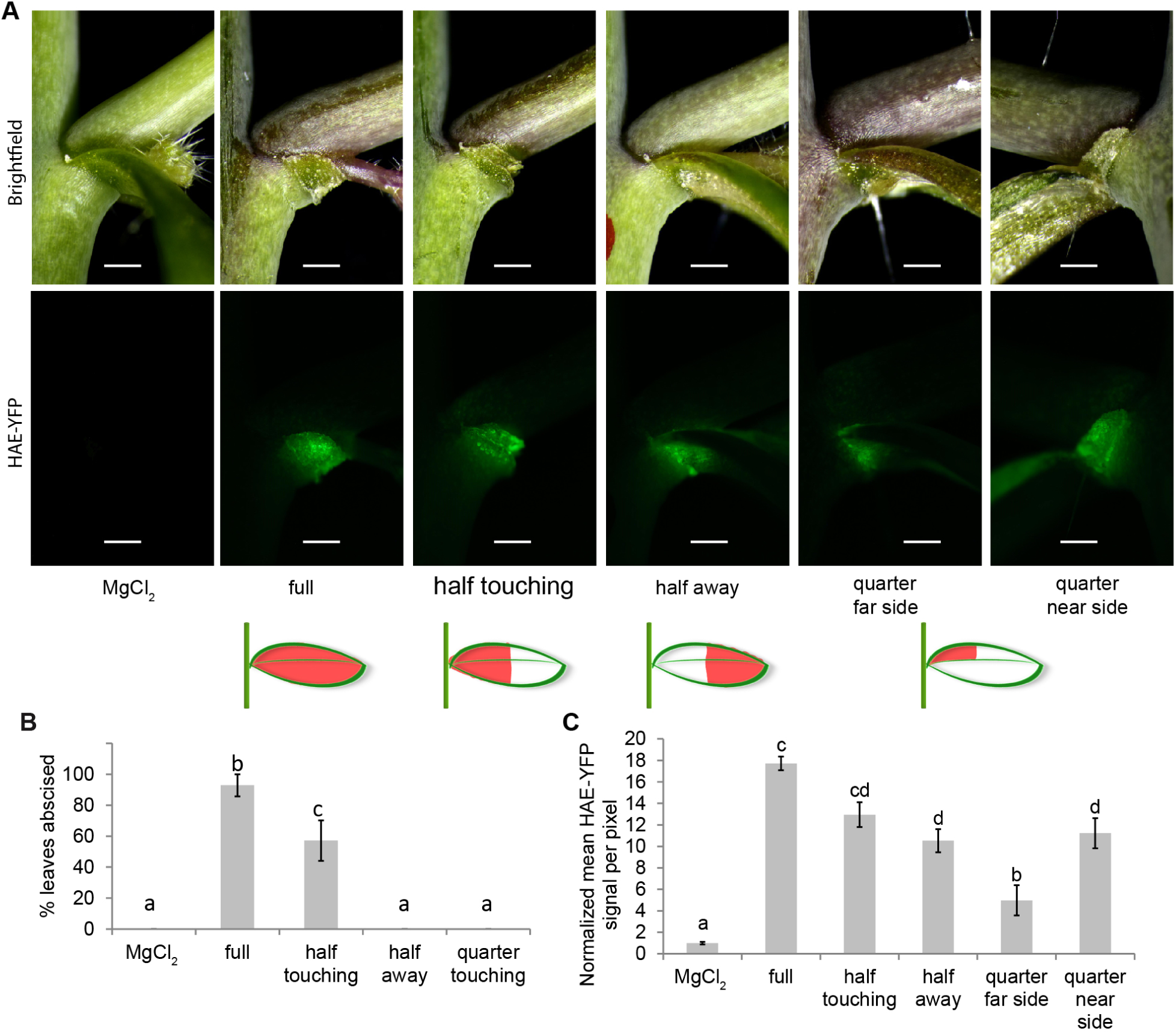
Abscission occurs when DC3000 touches the AZ, however, long distance signals are also sent from distal portions of the leaf to the AZ. (A) Leaves infiltrated in indicated portions of the leaf shown 3 days after infection. The same samples are shown in the top and bottom panels where the top panels are imaged with reflected white light and the bottom panels are imaging YFP florescence. (B) Percent of cauline leaves to abscise three days after treatment. Data are mean ± s.e.m.; n=7 biological replicates (one plant each); letters indicated different statistical quantities t-test *P* < 0.05. (C) Quantification of HAE-YFP fluorescence two days after infection while all leaves were still attached. Data are mean ± s.e.m.; n=4 biological replicates (one plant each); letters indicate different statistical quantities t-test *P* < 0.05. Scale bar is 0.5 mm.

### A signal is transduced across the uninfected half of the cauline leaf to the AZ in response to distal bacterial infection

To differentiate between the two possibilities of either bacteria moving or plants signaling to the AZ remotely, we designed an experiment to test if DC3000 moved in cauline leaves. DC3000 carrying the luciferase gene driven by the constitutive kanamycin promoter was infiltrated into the distal half (away from the AZ) of cauline leaves while the proximal half of the leaf was not infiltrated. A mark was drawn on the leaf to mark the boundary between infiltrated and not infiltrated. Two days after infection the cauline leaves were removed and cut in half along the boundary mark. Bacteria remained exclusively where they were infiltrated as the uninfected leaf half did not have detectable luciferase signal (Figure 4). This result indicates that there must be a signal transduced across the uninfected portion of the leaf that triggers AZ cell swelling, partial cell separation, and expression of HAE-YFP.

**Figure 4.**
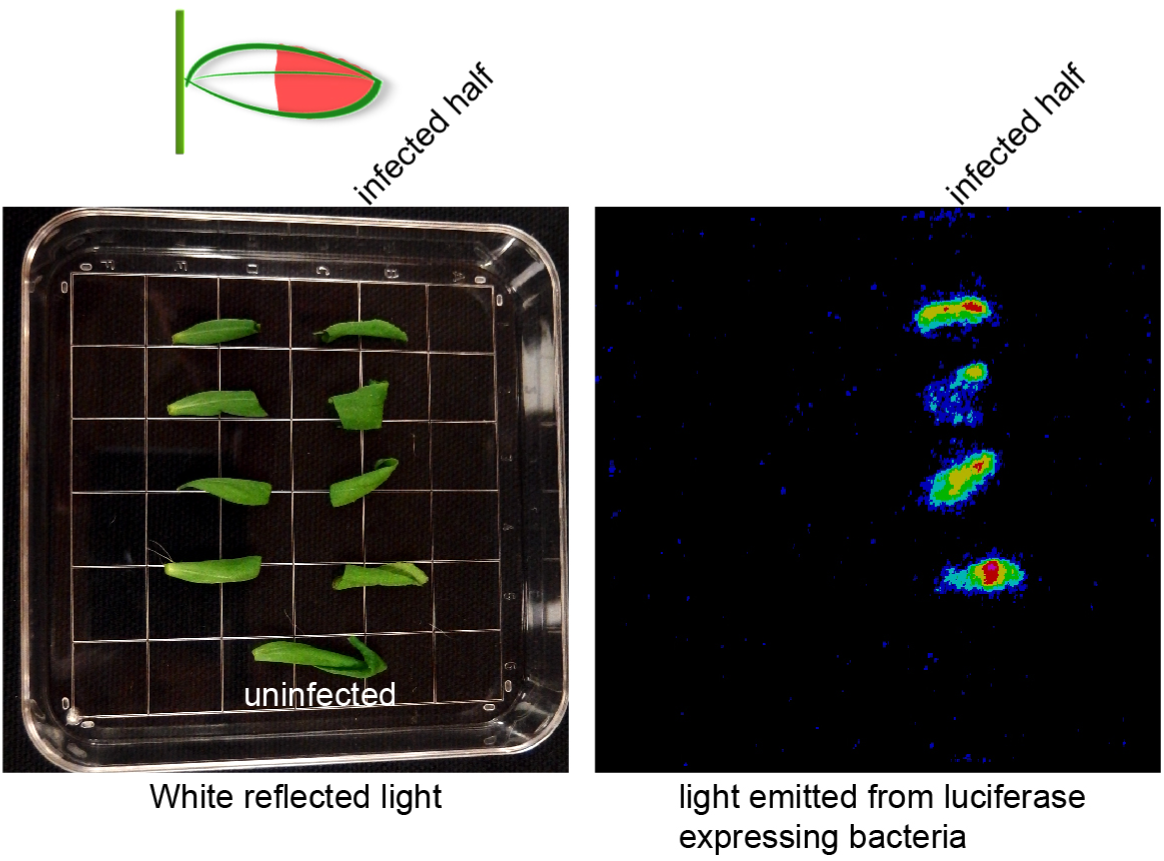
DC3000 does not move in cauline leaves from the place of infiltration. The indicated half of each cauline leaf was infiltrated with DC3000 that express luciferase constitutively driven by the kanamycin promoter. A line was drawn on the leaf with a pen to indicate the border of the infiltrated/not infiltrated. The leaves where cut in half 2 days after infection and imaged with white reflected light (left) and luminescence (right). Four replicates were performed with the same results.

### The floral organ abscission pathway is required for DC3000 triggered cauline leaf abscission

Previous work has shown that much of the floral organ abscission signaling pathway is conserved in the pathway for drought-triggered leaf abscission [8]. We hypothesized that pathogen-triggered leaf abscission would also require the core floral abscission signaling pathway. To test this hypothesis, floral organ abscission defective mutants were treated with DC3000 and AZ morphology and abscission were scored (Figure 5). *hae hsl2, ida*, and *nev* mutants all had statistically reduced abscission after pathogen treatment (Figure 5B). *hae hsl2* and *nev* mutants were completely blocked in abscission while *ida* had quantitatively reduced leaf abscission (Figure 5B). Interestingly, while *hae hsl2* mutants could not shed their leaves, their AZ cells did enlarge from the DC3000 treatment (Figure 5C). This suggests that AZ cell enlargement does not require *hae hsl2* (or *ida*). The ability to uncouple AZ cell enlargement from cell separation suggests that AZ cell enlargement and cell separation are distinct phases of abscission. It is interesting that Arabidopsis possesses a master cell separation signaling pathway that governs all known abscission events. It will be equally interesting to see if this core abscission pathway extends beyond Arabidopsis.

**Figure 5.**
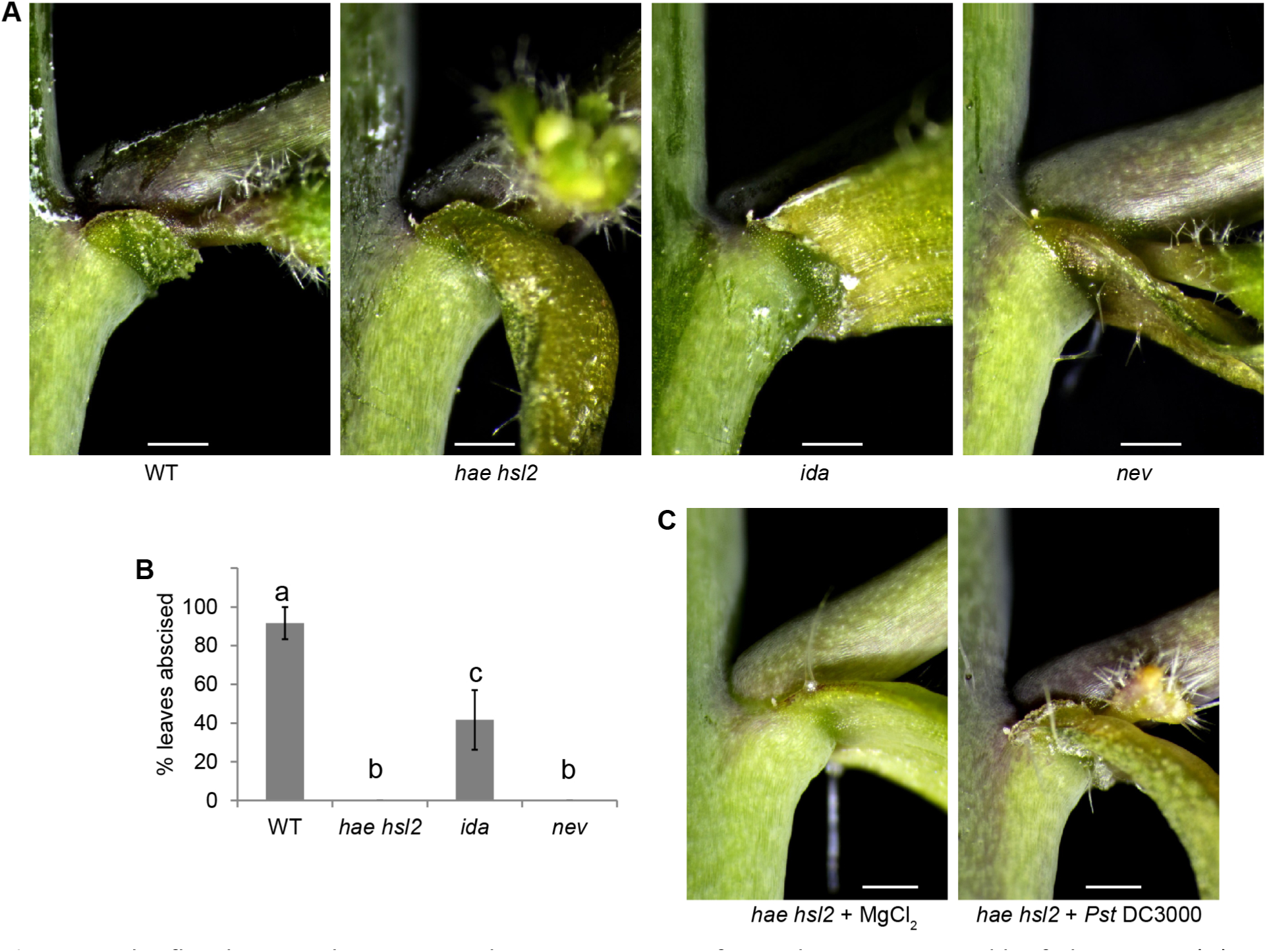
The floral organ abscission pathway is necessary for pathogen-triggered leaf abscission. (A) Photos of cauline leaf AZ of WT plants and floral abscission defective mutants treated with DC3000 for 3 days. (B) Percent cauline leaves abscised 3 days after infection. Data are mean ± s.e.m.; n=6 biological replicates (one plant each); letters indicated different statistical quantities t-test *P* < 0.05. (C) Micrograph of cauline leaf AZs from *hae hsl2* plants treated with or without DC3000. Images are representative from at least 4 replicates. Scale bar is 0.5 mm.

### Mutants with impaired bacterial defense fail to shed their leaves normally in response to DC3000 treatment

As shown, *PAD4* and *EDS1* are co-expressed with *HAE* (Figure 1). Also, *HAE* expression appears to be correlated with SA levels. Additionally, HAE appears to share the same co-receptor, BRI1-ASSOCIATED RECEPTOR KINASE (BAK1), as the receptor for perceiving bacterial flagellin, FLAGELLIN-SENSITIVE 2 (FLS2) [12]. Therefore, we asked whether several genes necessary for bacterial defense might also play a role in abscission. We found that plants with reduced levels of SA, NahG and *sid2*, had quantitatively reduced abscission, with NahG being essentially qualitatively blocked in abscission (Figure 6A). Additionally, *pad4, eds1*, and *pad4 sag101* (*SENESCENCE-ASSOCIATED GENE 101*) mutants also had quantitatively impaired abscission. *pad4* was more severely impaired in abscission than *eds1*, while the double mutant *pad4 sag101* had a slightly more severe phenotype that *pad4* alone (Figure 6A). These results strongly suggest that DC3000 triggered leaf abscission is a bona fide defense response. On the other hand, if leaves that were damaged or sick simply fell off passively, we would expect defense mutants to shed their leaves more readily than WT since the mutant leaves bear more disease symptoms than WT.

**Figure 6.**
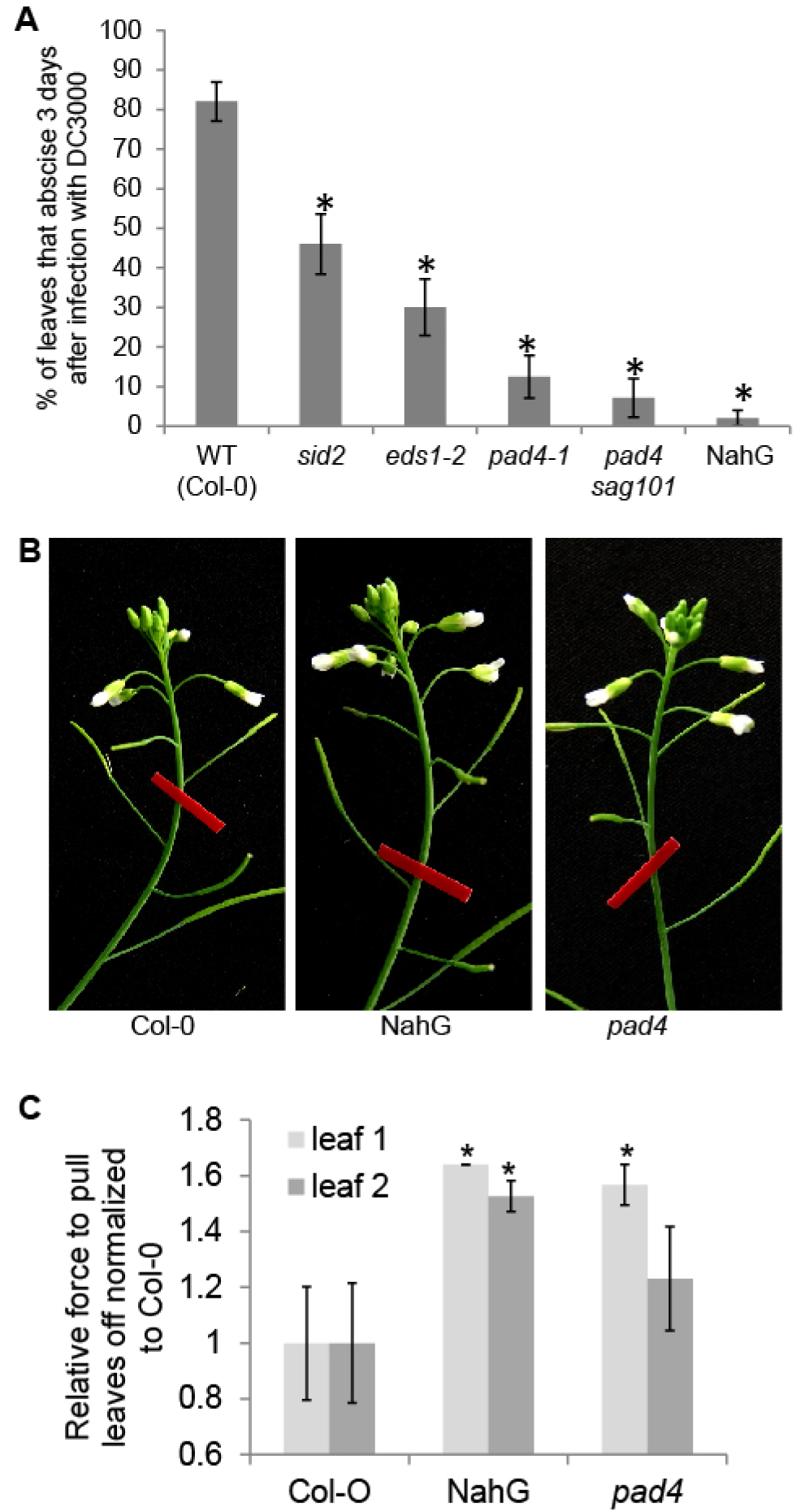
Pathogen defense defective mutants are defective in pathogen-triggered cauline leaf abscission. (A) Percent of cauline leaves that abscised 3 days after infection. Data are mean ± s.e.m.; n=25 biological replicates (one plant each); t-test versus WT; * *P* < 0.0001. (B) NahG and *pad4* floral organ abscission is similar to WT. Red tape is 9.5 mm wide. (C) Relative leaf breakstrength force of WT (Col-0), NahG transgenic plants, and *pad4* exposed to drought and re-watering conditions. Data are mean ± s.e.m.; n=12 biological replicates (one plant each); t-test versus WT; * *P* < 0.05.

As mentioned, *SID2* (*ICS1*), *ICS2, PAD4*, and *EDS1* are all transcriptionally up-regulated in stamen abscission zones through the process of floral organ abscission [27]. Therefore, we addressed whether these genes may also be necessary for floral organ abscission. Neither NahG (SA deficient) nor *pad4* plants had obviously different floral organ abscission (Figure 6B). This represents a major difference between floral organ abscission and pathogen-triggered leaf abscission. Next we asked if NahG trangenic plants or *pad4* mutants had altered drought-triggered cauline leaf abscission. Surprisingly, NahG transgenic plants had statistically reduced abscission in cauline leaves 1 and 2 while *pad4* had reduced abscission in cauline leaf 1 (Figure 6C). This suggests SA or SA signaling plays a role in drought-triggered cauline leaf abscission.

Plant’s defense response to bacterial pathogens has largely been studied in Arabidopsis tissue that is fairly vulnerable to bacterial colonization. The preferred system of study is four week old rosette leaves from plants that are not flowering (grown with 8-12 hrs light per day). To further assist bacterial colonization, plants are typically grown in 70-90% relative humidity. The rosette leaves from nonflowering plants, grown under high humidity, can support up to 5 logs of growth in 3 days [26]. Our cauline leaf system has numerous differences from the typical Arabidopsis pathosystem. Our system uses cauline leaves from flowering plants grown in long days (16 hrs light per day) at 50-65% relative humidity. Bacterial enumeration experiments were performed to assess the level of bacterial colonization in our abscission system. In general, mutants known to be defective in defense allow more bacterial enumeration than does WT (Figure 7). In two days DC3000 had enumerated slightly less than 2 logs in WT while it enumerated 2.5-3 logs in Arabidopsis defense mutants (Figure 7). *sid2* mutants were the least compromised mutant tested in terms of both bacterial enumeration and cauline leaf abscission. While *SID2*, also known as *ISOCHORISMATE SYNTHASE 1* (*ICS1*), is the major limiting gene in the synthesis of SA in rosette leaves from non-flowering plants, *ICS2* may function redundantly in tissues on flowering plants. In fact, *ICS2* is induced in stamen abscission zone through the process of abscission [27]. Overall, cauline leaves, grown in our standard conditions, allowed less bacterial multiplication in WT or defense mutants than did rosette leaves from non-flowering plants, but displayed the same relative levels of susceptibility.

**Figure 7.**
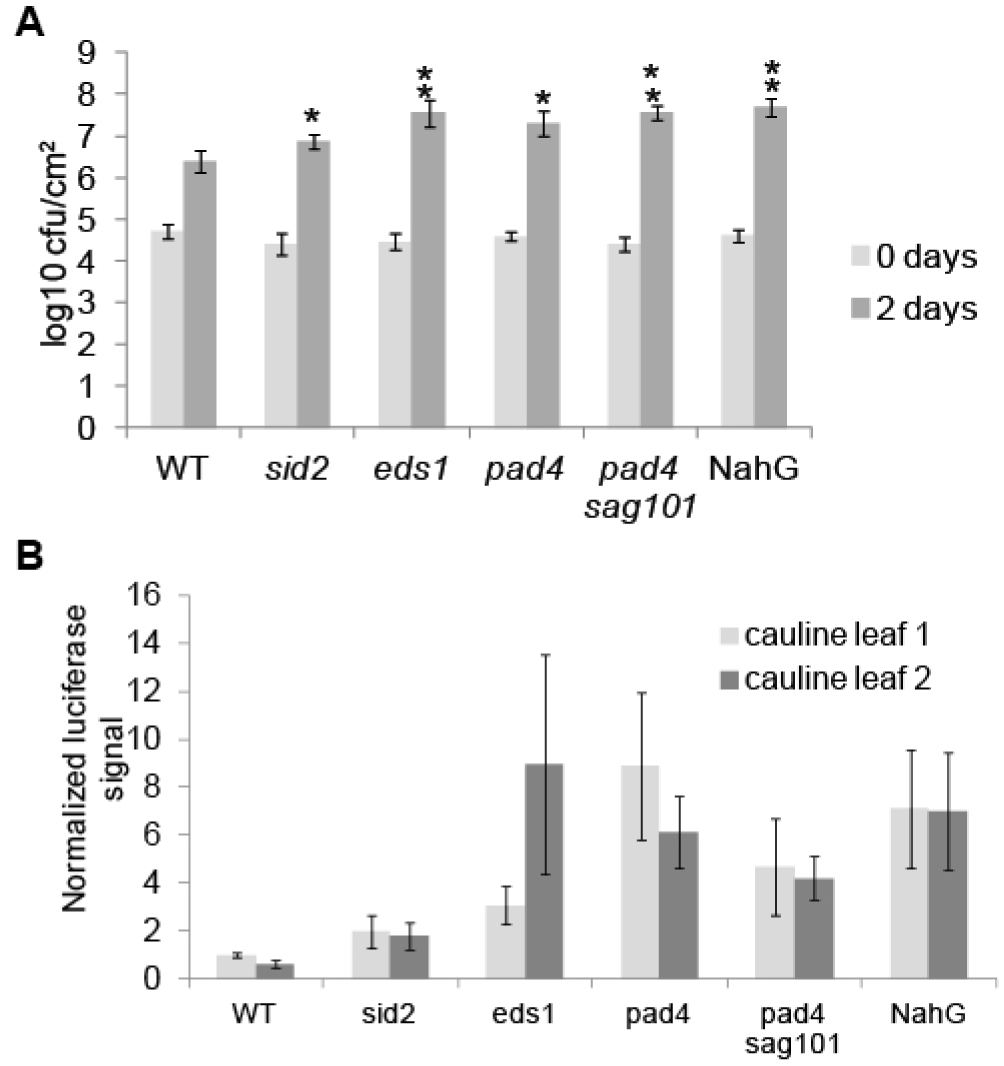
Bacterial growth in cauline leaves of Arabidopsis defense mutants. (A) Bacterial enumeration of cauline leaf 2. Data are mean ± s.e.m.; n=5 biological replicates (one plant each); t-test versus WT; * *P* < 0.1, ** *P* < 0.05. (B) Luciferase signal from plants infected for 2 days with O.D. = 0.001 DC3000 lux. Data are mean ± s.e.m.; n=5 biological replicates (one plant each).

## Discussion

Resistance protein-mediated effector-triggered immunity involves the host recognizing that something has changed within itself due to pathogens trying to suppress the host’s immune system. Once the host recognizes these changes, a strong defense response is launched. In plants, this strong defense response sometimes takes the form of the hypersensitive response in which the infected tissue is intentionally destroyed to limit further pathogen colonization. Here we show a form of defense triggered by a bacterial effector or effectors that differs from the classical resistance protein-based ETI response in that it is launched at the organ level by a virulent pathogen. Leaves that have a functional AZ can simply be shed to limit further pathogen colonization and alleviate the need for further resource consumption. Essentially, plants can cut their losses since they can always produce more leaves. Importantly, shedding infected leaves eliminates 100% of the bacteria in the infected leaf from the plant body. In contrast, many bacterial growth controlling plant defense mechanisms only result in 10-1000 fold reduction of bacterial growth. *Pseudomonas syringae* spreads by raindrop momentum, wind, and insects [22–25,35]. Therefore, eliminating disease source leaves reduces the possibility of infection spreading to healthy tissue. One could imagine that abscising leaves infected with pathogens that can spread systemically throughout the plant could be even more beneficial to the plant.

Pathogen-triggered leaf abscission appears to be an effector-triggered response and not a PTI response since DC3000 without a functional type III secretion system cannot trigger leaf abscission. This finding also excludes the bacterial toxin coronatine as the trigger of abscission. It is not clear at the moment which effector(s) trigger leaf abscission. Both DC3000 with and without effectors *avrRpm1* and *avrRps4* trigger leaf abscission. These findings cannot exclude the possibility that AvrRpm1 and AvrRps4 can trigger leaf abscission by themselves. However, it does indicate an endogenous effector of DC3000, that is injected into the plant via the type III secretion system, can trigger leaf abscission. Pathogen-triggered leaf abscission is a relatively slow response in comparison to the hypersensitive response, which can occur in as little as 8 hours after infection. However, leaf abscission will of course ultimately supersede all other defense responses in the leaf.

The defense response to DC3000 in cauline leaves activates HAE expression and requires the floral organ abscission signaling pathway for abscission to occur. Previously, HAE has been proposed to be a positive regulator of bacterial growth in rosette leaves, where HAE appears to work on the bacteria’s behalf in degrading pectin to facilitate bacterial colonization [15]. We propose a broad function of HAE may be to determine the sacrifice of tissues or organs that are infected so that the rest of the plant can live. Currently, no known condition triggers rosette leaf abscission in Arabidopsis. However, rosette leaves do senesce and HAE may function in initiating this process once bacterial titers in the infected leaf have surpassed a threshold. In support of this hypothesis, sepals in *hae hsl2* plants senesce later than WT sepals do, which suggests HAE/HSL2 may have a part to play in senescence.

The abscission process can be divided into several phases. Floral organ abscission has been divided into 4 phases. In phase 1 abscission zones develop. During phase 2 abscission zones become competent for abscission. In phase 3 cell separation is initiated. Finally in phase 4 abscission zone cells become enlarged and differentiated [36]. Interestingly, in pathogen-triggered leaf abscission *hae hsl2* mutants are only deficient in cell separation. Abscission zone cells clearly become enlarged after infection with DC3000, however, they do not abscise. Furthermore, AZ cells begin enlarging in WT plants at two days after infection, which is one day before leaves abscise. It is not clear if the phases of abscission are ordered slightly differently in leaf abscission and floral organ abscission. It is also possible that AZ cell enlargement in floral organ AZs begins before cell separation and continues after cell separation. Floral organ AZs cannot be visualized nondestructively prior to abscission because sepals and petals cover the AZ. In contrast, cauline leaf abscission zones are ideal for real time monitoring since they are not obscured by other tissues. Detailed physiological and anatomical measurements of the three Arabidopsis abscission systems (floral organ, drought‐ and pathogen-triggered leaf abscission) over time may shed light on the order of the phases and the actual function of AZ cell enlargement.

Pathogen-triggered leaf abscission appears to be an active defense response that requires components needed for rosette leaf defense. The defense mutants *pad4, eds1, sid2, pad4 sag101*, and NahG transgenic plants all fail to abscise normally after infection with DC3000. If leaf abscission were occurring simply because leaves were sick and damaged, the expectation would be that the defense mutants would abscise more readily than WT. There is evidence that abscission components and defense components are physically associated in protein complexes. HAE physically interacts with BAK1 where BAK1 serves as a co-receptor for HAE [12]. BAK1 is also the co-receptor for FLS2 which perceives bacterial flagellin [37]. HAE physically interacts with CST which physically interacts with EVR [38]. *EVR* is also known as *SUPPRESSOR OF BIR1 1* (*SOBIR1*) because mutations in it can suppress *bak1-interacting receptor-like kinase 1* (*bir1*), which has a constitutive defense response phenotype [39]. Overactive defense responses in *bir1* partially require *EDS1* and *PAD4* [39]. Unfortunately, none of the above interactions have been demonstrated in AZs. Instead, the above protein-protein interactions with HAE have been demonstrated in mesophyll protoplasts. There appear to be many future opportunities to use cross reference analysis to further both the defense and abscission fields.

Salicylic acid production or signaling appears to be required for proper pathogen-triggered leaf abscission to occur. Transgenic plants over-expressing NahG are almost entirely blocked in leaf abscission. *sid2* (or *ics1*) also have quantitatively reduced leaf abscission. Differences in severity of the abscission defect between NahG and *sid2* could be explained by alternate means of producing SA in cauline leaves of *sid2* plants. *ICS2* is transcriptionally induced through the process of floral organ abscission and may also be expressed in cauline leaves [27]. On the other hand, NahG plants are likely to have a more uniform reduction of SA throughout the entire plant than *sid2. sid2* plants have a relatively mild defect in restricting bacterial growth in cauline leaves compared to *eds1, pad4, pad4 sag101*, and NahG, which supports the idea that redundant methods of producing SA may be present in *sid2* plants. *pad4* and *eds1* mutants also accumulate less SA as they are thought to signal activation of ICS1 by transducing reactive oxygen species signals and also participate in a positive feedback loop of SA amplification [40–42]. SA is not required for developmentally timed floral organ abscission because NahG plants abscise their floral organs normally. However, in an unexpected twist, SA influences drought-triggered leaf abscission. This infers there may be a tight connection between senescence and abscission in leaves. We have never observed completely green leaves abscising after drought treatment [8]. Instead, leaves always turn at least partially yellow before abscising. While SA’s role in leaf abscission has not been well characterized in the literature, the gaseous hormone ethylene has been implicated in pathogen-triggered leaf abscission in tomato, pepper, and peanut [16,18,21]. Ethylene would likely regulate pathogen-triggered leaf abscission in Arabidopsis since ethylene insensitive plants have already been shown to have delayed floral organ abscission and are also more tolerant to *Pseudomonas syringae* [43,44].

The location of a bacterial infection on a leaf determines the extent of abscission. Abscission occurs when the bacterial infection is in the base of the cauline leaf that touches the AZ. However, only partial cell separation occurs when a portion of the leaf is infected that is distal to the abscission zone. One possible reason for this response could be that it prevents spread of the infection to the rest of the plant; however, DC3000 is not actually mobile in cauline leaves. Perhaps this response occurs as a general response in case the pathogen is mobile. Alternatively, abscission might simply occur because once the base of the cauline leaf is severely compromised, the distal portion of the leaf would not be able to survive. It is not clear what mobile signal could be transducing a signal from the distal infected leaf tissue, across non-infected tissue, to the AZ. Potentially this system could be an attractive model system for studying cell to cell communication. Compared to proximally infiltrated leaves, fully infiltrated leaves abscised at a higher percentage that was statistically significant. Therefore, remote signaling from the distal portion of the cauline leaf increases abscission. Remote signaling of infection could also provide the AZ an early warning in the case of mobile pathogens.

In conclusion, we define a new model system for studying pathogen-triggered leaf abscission. Our study begins to explain the genetics governing pathogen-triggered leaf abscission. The abscission pathway first found to regulate floral abscission is required for all known forms of inducible leaf abscission in Arabidopsis. We found that the previously disparate pathways regulating defense and abscission are connected so that a number of defense components are necessary for pathogen-triggered leaf abscission. Additionally, salicylic acid is not only necessary for full pathogen-triggered leaf abscission but also drought-triggered leaf abscission. We propose a threshold model of defense for cauline leaves where cauline leaves attempt to fight minor infections until the threat is too great and abscission is activated (Figure 8).

**Figure 8.**
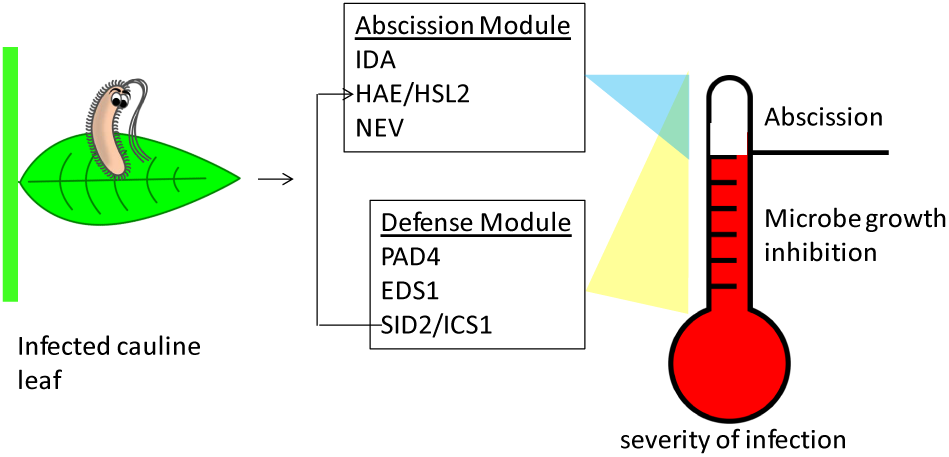
Proposed cauline leaf defense model. Cauline leaf microbial defense has a threshold system. Minor infections are fought to limit the multiplication of bacterial growth. Microbial growth inhibition requires *PAD4, EDS1, SAG101*, and *SID2* (and salicylic acid). If the infection becomes too serious, the entire cauline leaf will simply be abscised. Abscission requires the previously mentioned defense components as well as *HAE/HSL2, IDA*, and *NEV*. The defense module potentially regulates the abscission module via salicylic acid where salicylic acid induces expression of *HAE*.

## Material and Methods

### Plant material and growth conditions

The Columbia accession of Arabidopsis was used as a wild type (Col-0; ABRC stock# CS70000). Mutants used were the indicated allele: *hae-3 hsl2-3, ida-2, nev-3, pad4-1, eds1-2, sid2-2, sag101-2 pad4-1* [32,9,45–47,29,48]. Plants carrying HAE-YFP driven by the native promoter were previously described [8]. Plants were grown in Promix BX (Premier Tech Horticulture) at 23°C, 16 h light / 8 h dark, 100-150 μE·m^-2^·s^-1^, and 50-70% humidity (except for indicated experiments with 8 h light / 16 h dark, >75% humidity). Plants were planted in a randomized complete block experimental design.

### Co-expression analysis

Publicly available microarray data was downloaded from Gene Expression Omnibus (GSE39385, GSE5727, GSE19255, GSE10646, GSE5632) and ArrayExpress (E-MEXP-173, E-MEXP-1474) [49,50,31,51,27,52] and reanalyzed with RobiNA using the PLIER algorithm [53].

### Bacterial induced cauline leaf abscission

Various strains of *Pseudomonas syringae* pv. tomato strain DC3000 were grown on King’s B plates for 2-3 days at room temperature. Needless syringe infiltration of leaves was performed by scraping bacteria off of 2-3 day old plates and resuspending them in 10 mM MgCl_2_ to an A_600_ = 0.01 (unless otherwise indicated) for standard abscission assays. DC3000 COR^-^ is CFA^-^ CMA^-^ [34]. The first two cauline leaves were infiltrated on each plant. Three days after infection cauline leaves 1 and 2 were gently touched to see if abscission had occurred. Abscission per plant was scored as 0%, 50%, or 100% depending if 0, 1, or 2 cauline leaves abscised (out of two possible). Cauline leaf breakstrength was measured as previously described [8,54].

### Pseudomonas growth measurements

Bacterial enumeration assays were performed by infiltrating leaves with bacteria at an A_600_ = 0.0005. Bacterial titer in leaf punches was determined as previously described [26]. Quantification of luminescent bacteria was performed by infiltrating leaves with DC3000 that expresses luciferase driven by the constitutive kanamycin promoter (LuxCDABE operon) at a concentration of A_600_ = 0.001 [55]. Luminescence from bacterial luciferase in leaves was quantified two days after infection with a HRPCS4 photon-detection camera and IFS32 software (Photek Ltd, http://www.photek.com) where data was integrated for 120 seconds.

### Microscopy

Fluorescent and brightfield microscopy was performed as previously described [8]. Brightfield images are extended depth of field images and YPF images are a single depth of field. Quantification of HAE-YFP signal was performed with ImageJ where mean pixel intensity was calculated for AZs two days after infection.

### Assay of DC3000 movement within cauline leaves

DC3000 lux at a concentration of A_600_ = 0.001 was infiltrated into the distal half of cauline leaves. A line was drawn on the cauline leaves to indicate the boundary between infiltrated and uninfiltrated portions. Two days after infection, cauline leaves were removed and cut in half along the boundary mark and luciferase luminescence was imaged with a HRPCS4 photon-detection camera (described above).

## Author Contributions

O.R.P. conceived the study, O.R.P and W.G. designed the experiments, O.R.P. performed and analyzed the experiments, O.R.P, W.G., and J.C.W. wrote the paper.

## Acknowledgements

We thank Catherine Espinoza for reading and editing the manuscript, Scott Peck for the use of his photon-detection camera used to quantify bacterial luciferase, and Barbara Kunkel for providing the coronatine mutant bacterial strains.

---

**Supplemental Figure 1.**
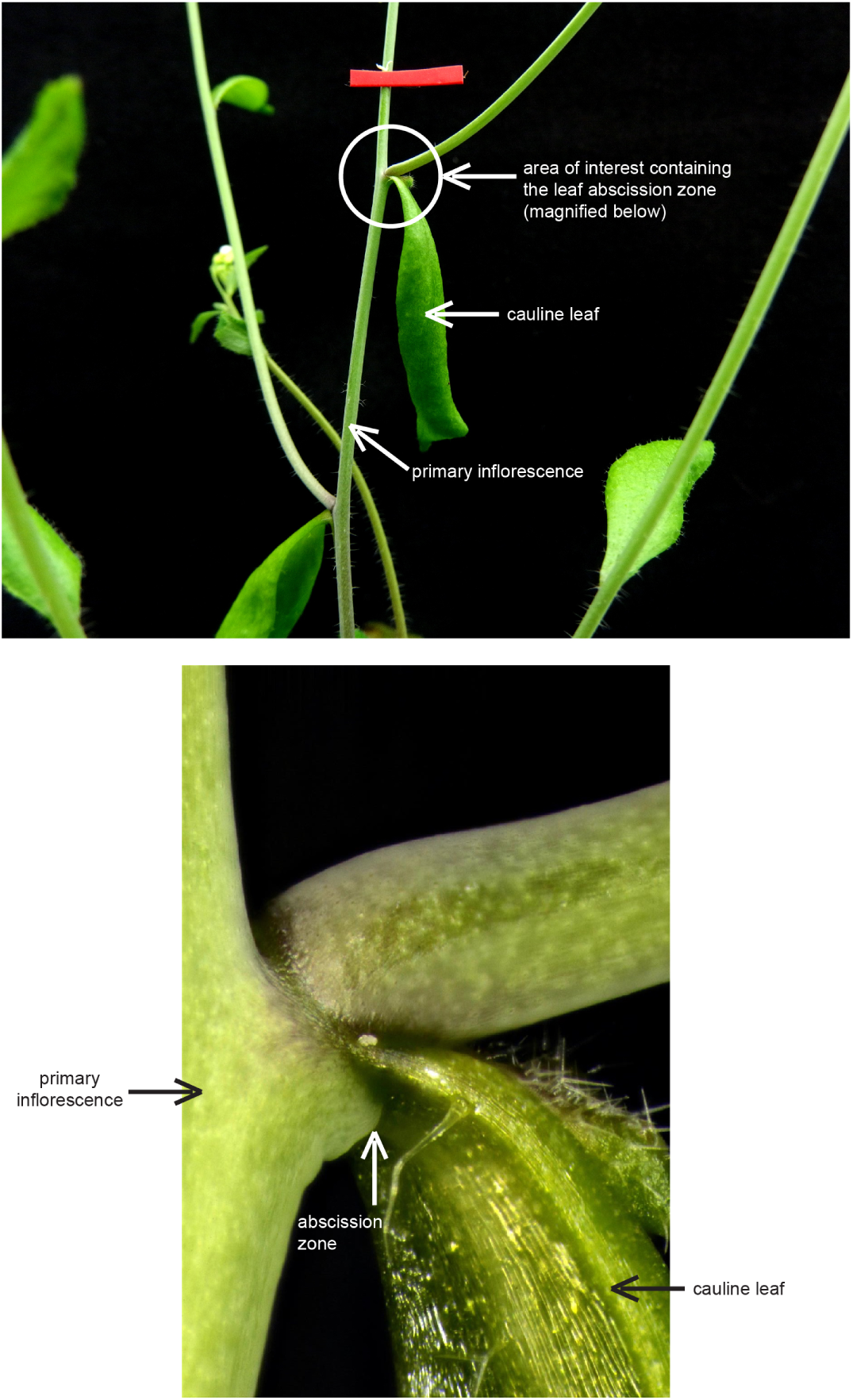
The cauline leaf abscission system. The top panel shows the second cauline leaf on the primary inflorescence. The bottom panel is a magnification from the circled area in the top panel. The cauline leaf abscission zone enables cauline leaves to be shed.

**Supplemental Figure 2.**
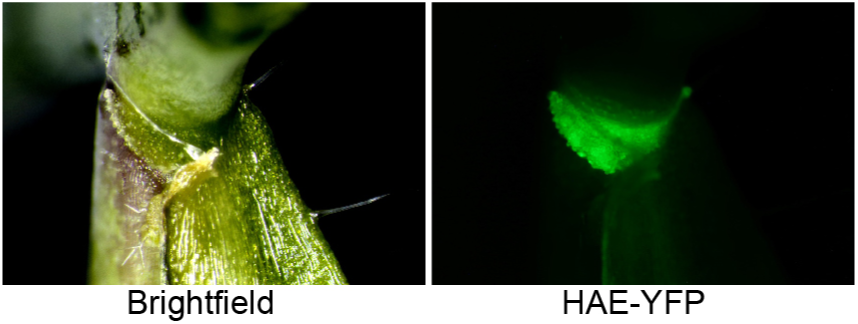
View from the bottom of a cauline leaf infected on the left quarter which is touching the AZ. Note the left side of the cauline leaf has peeled off of the abscission zone while the right side of the cauline leaf remains attached.

**Supplemental Figure 3.**
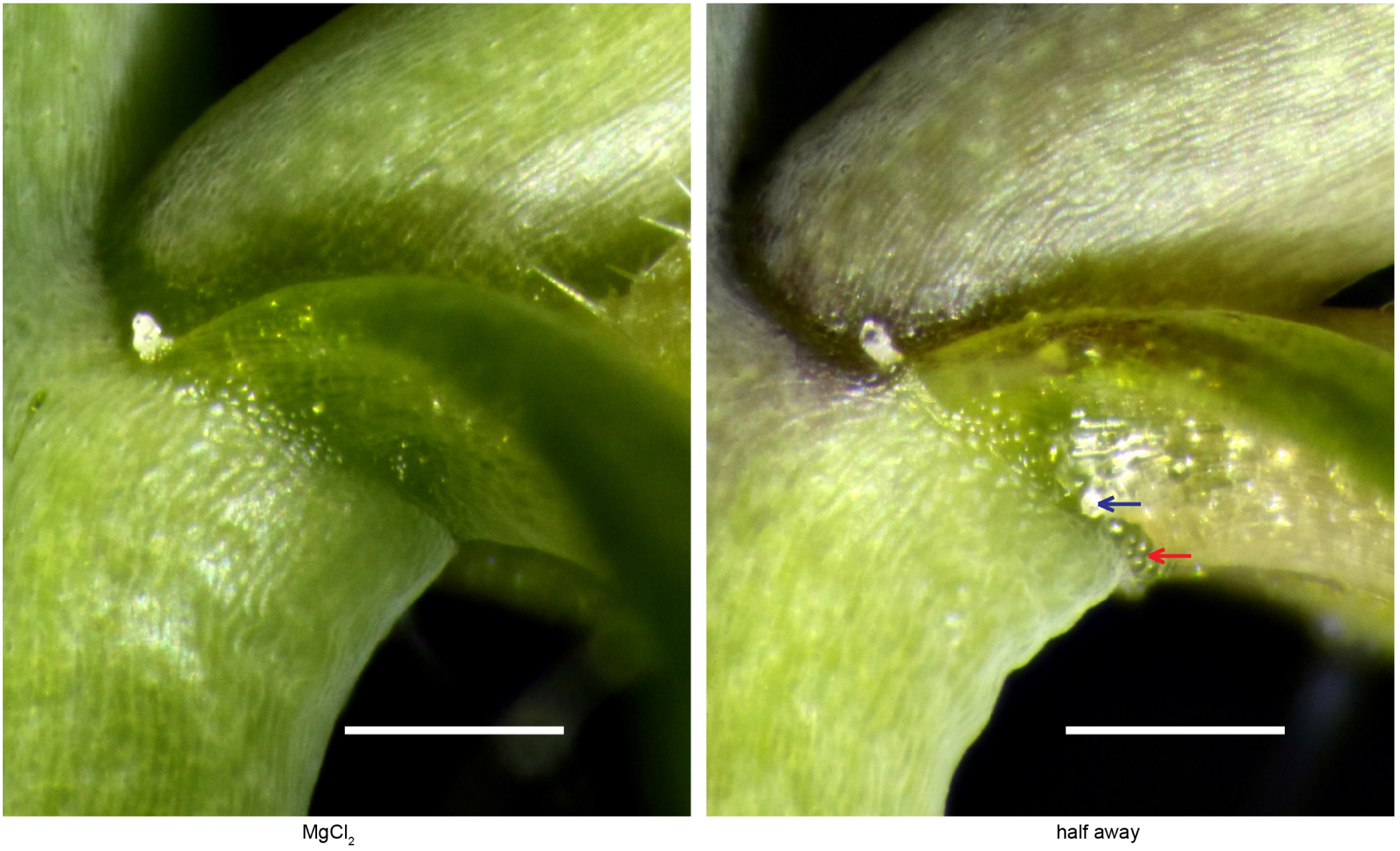
Enlargement of control MgCl_2_ treatment and distal half DC3000 infection (half away) from figure 3. Blue arrow indicates area where cell separation has occurred. Red arrow indicates swollen abscission zone cells.

**Supplemental Figure 4.**
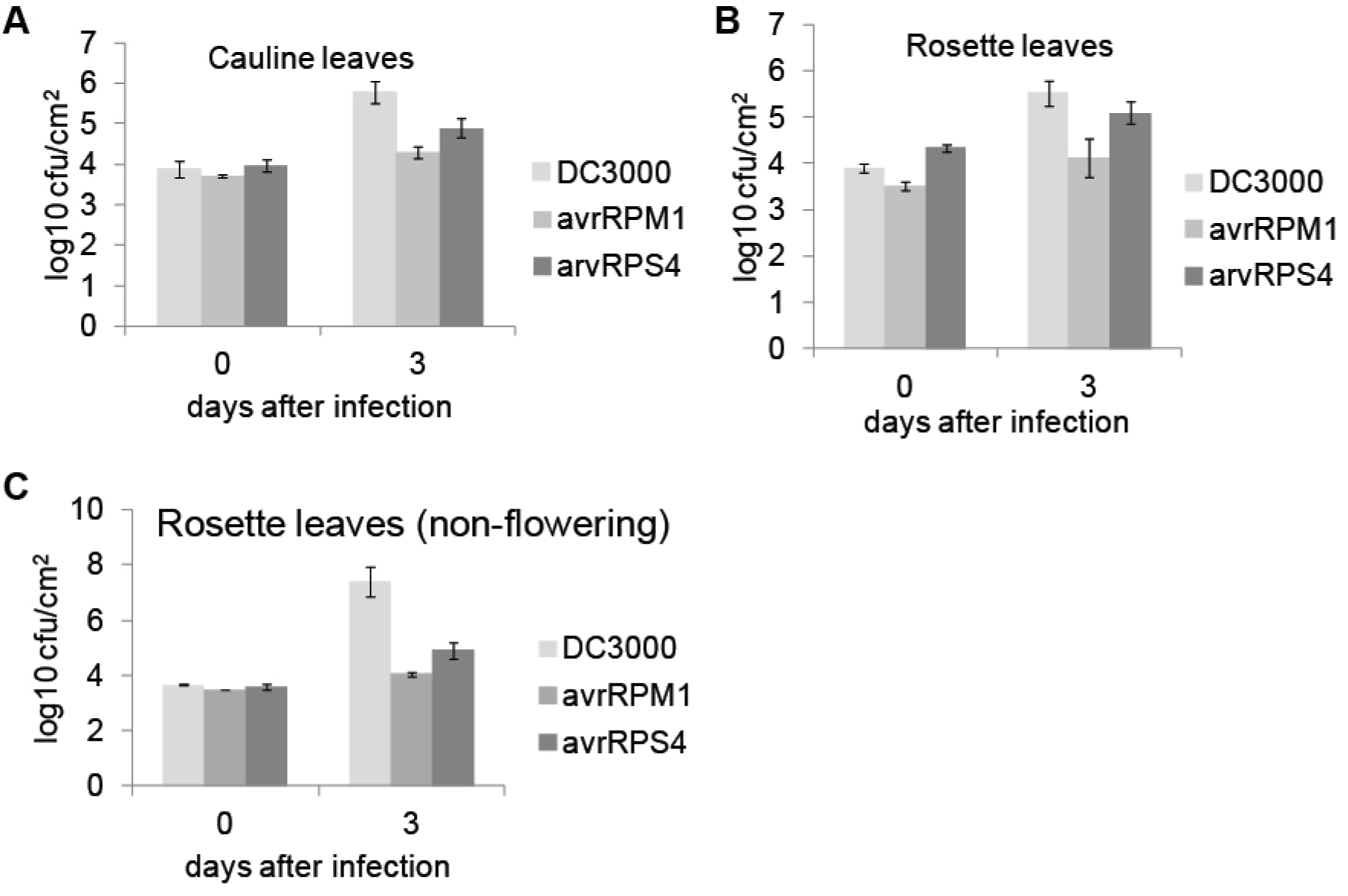
Flowering plants are more resistant to *Pst* than non-flowering plants. Bacterial enumeration in (A) cauline leaves and (B) rosette leaves from flowering plants grown in 16 h light / 8 h dark with 50-65% relative humidity. Bacterial enumeration of non-flowering plants grown in 8 h light / 16 h dark with ≥ 75% relative humidity.

